# Genetic evidence for environment-dependent hybrid incompatibilities in threespine stickleback

**DOI:** 10.1101/2021.06.24.449805

**Authors:** Ken A. Thompson, Catherine L. Peichel, Diana J. Rennison, Matthew D. McGee, Arianne Y. K. Albert, Timothy H. Vines, Anna K. Greenwood, Abigail R. Wark, Yaniv Brandvain, Molly Schumer, Dolph Schluter

## Abstract

Hybrid incompatibilities occur when interactions between opposite-ancestry alleles at different loci reduce the fitness of hybrids. Most work on incompatibilities has focused on those that are ‘intrinsic’, meaning they affect viability and sterility in the laboratory. Theory predicts that ecological selection can also underlie hybrid incompatibilities, but tests of this hypothesis are scarce. In this article, we compiled genetic data for F_2_ hybrid crosses between divergent populations of threespine stickleback fish (*Gasterosteus aculeatus* L.) that were born and raised in either the field (semi-natural experimental ponds) or the laboratory (aquaria). We tested for differences in excess heterozygosity between these two environments at ancestry informative loci—a genetic signature of selection against incompatibilities. We found that excess ancestry heterozygosity was elevated by approximately 3% in crosses raised in ponds compared to those raised in aquaria. Previous results from F1 hybrids in the field suggest that pond-specific (single-locus) heterosis is unlikely to explain this finding. Our study suggests that, in stickleback, a coarse signal of environment-dependent hybrid incompatibilities is reliably detectable and that extrinsic incompatibilities have evolved before intrinsic incompatibilities.

---

Hybrid incompatibilities—interactions among divergent genetic loci that reduce the fitness of hybrids—are a key component of reproductive isolation between diverging lineages (Coyne and Orr 2004). Incompatibilities have been studied most intensively in the context of sterility and mortality, in part because these traits are conducive to reliable phenotyping in the laboratory and can be underpinned by few loci (Fishman and Sweigart 2018; Maheshwari and Barbash 2011). These sorts of barriers have come to be called ‘intrinsic’ hybrid incompatibilities due to the fact that there are conflicts within the hybrid genome that are expected to impact hybrids in most environmental contexts (though note that the strength of selection against some intrinsic incompatibilities can vary across environments [Demuth and Wade 2007]). Studies have shown that the number of intrinsic in-compatibilities tends to increase with genetic divergence between parents (Matute et al. 2010; Moyle and Nakazato 2010; Wang et al. 2015) and that incompatibilities can be common throughout the genomes of isolated conspecific populations (Corbett-Detig et al., 2013). Collectively, evolutionary biologists have made substantial progress toward identifying generalities about the evolutionary genetics of intrinsic hybrid incompatibilities.

Ecological selection can underpin incompatibilities if particular allele combinations render hybrids unable to function in their ecological environment, such as in avoiding predators or capturing prey. Several recent studies have shown patterns consistent with this effect, wherein hybrids have ‘mismatched’ trait combinations and reduced fitness as a result (Arnegard et al., 2014; Thompson et al., 2021). Such studies have successfully demonstrated incompatibilities via interactions among traits, but links to the underlying genetics have not been made. Perhaps the most significant barrier to progress in studying the genetics of ‘ecological’ hybrid incompatibilities is the especial difficulty of detecting them. If many traits have a polygenic basis with small individual effect sizes (Rockman 2012), classic methods for detecting hybrid incompatibilities will be severely underpowered. For example, Arnegard et al. (2014) found that combining divergent jaw traits together in F_2_ threespine stickleback (*Gasterosteus aculeatus* L.) hybrids reduced their fitness because these traits interacted in a manner that reduced suction feeding performance. The interacting jaw components map to several regions of the genome that individually explain a small fraction (< 10%) of the phenotypic variance (and most variance was unexplained), thus rendering it difficult to study their individual epistatic fitness effects.

Recent theoretical advances, however, suggest ways to test for and measure the net effect of hybrid incompatibilities using experimental crosses. Specifically, selection against hybrid incompatibilities in an F_2_ hybrid cross causes an increase in ancestry heterozygosity—the number of sites in the genome that carry both parents’ alleles at ancestry-informative sites—at the population level (Barton and Gale, 1993; Simon et al., 2018). This is expected because F_2_ hybrids have a hybrid index of approximately 0.5—deriving half of their alleles from one parental species and half from the other. Thus, individuals with high heterozygosity relative to their hybrid index have fewer pairs of homozygous loci with opposite ancestry than relatively homozygous individuals with a similar hybrid index. Assuming most alleles affect the phenotype additively, having many loci with opposite homozygous ancestry can result in hybrids with maladaptive ‘mismatched’ phenotypes, whereas highly heterozygous individuals are expected to be relatively intermediate (Fig. 1 and Fig. S1). Whether ‘mismatch’ affects fitness, however, ultimately depends on the ecology of the system and the underlying fitness landscape. Such coarse approaches— coarse because they use summary statistics rather than direct mapping—are a promising means to identify the presence of small-effect hybrid incompatibilities at the genetic level using field experiments.

**Fig. 1.**
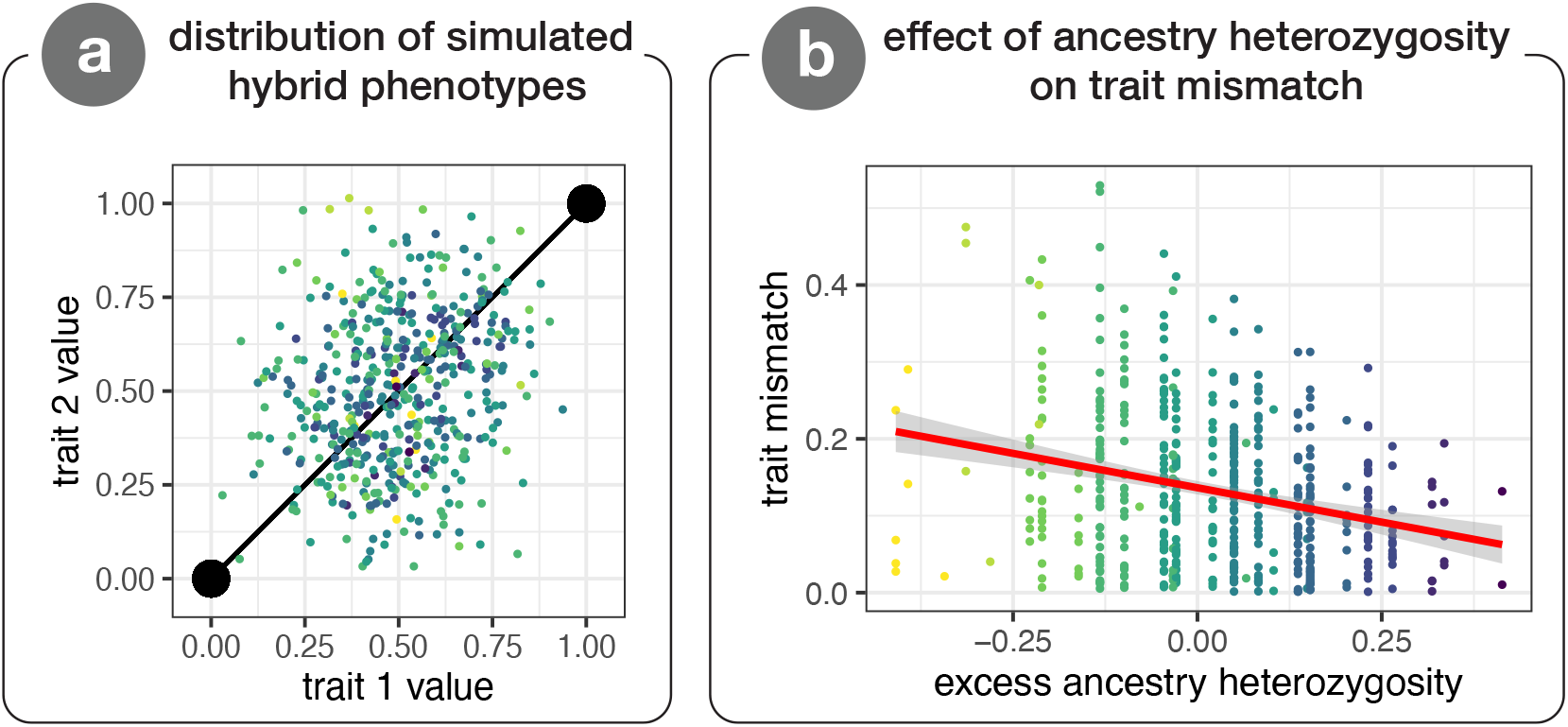
Results from simulations illustrating an ecological mechanism underlying the heterozygosity-incompatibility relationship in F_2_ hybrids. Both panels depict results from a representative simulation run of adaptive divergence and hybridization between two populations. We consider an organism with two traits that have both diverged as a result of selection. Coloured points are individual hybrids, with darker colours indicating higher heterozygosity. Panel (A) depicts the distribution of 500 F_2_ hybrid phenotypes in two-dimensional trait space. Large black points are the two parent phenotypes, which are connected by a black line indicating the ‘axis of divergence’. Panel (B) depicts the relationship between individual excess ancestry heterozygosity and trait ‘mismatch’ of individual hybrids (Thompson et al., 2021). Excess ancestry heterozygosity is is the observed heterozygosity minus the expected heterozygosity based on ancestry proportion—0 is the predicted mean in the absence of selection (approx. 0.5 observed heterozygosity). Mismatch is calculated as the shortest (i.e., perpendicular) distance between a hybrid’s phenotype and the black line connecting parents in (A). Variation parallel to this axis connecting parents in (A) captures variation in the ‘hybrid index’. The plot shows that trait mismatch is lower in more heterozygous F_2_ hybrids. Heterozygosity values are discrete because a small number of loci underlie adaptation in the plotted simulation run. See the Supplementary Methods for simulation methods and underlying assumptions. The ‘mismatch’–heterozygosity relationship is stronger, though less intuitive, in organisms with greater dimensionality (i.e., more traits; see Fig. S1 for a case with 10 traits following Barton 2001).

In this study, we compare patterns of selection on ancestry heterozygosity between F_2_ hybrid families raised in the lab to those from the same cross types raised in field enclosures. If ecological selection on ‘trait mismatch’ is operating in the field but not in the lab, selection for increased ancestry heterozygosity should be specific to the field (or at least stronger than in the lab). If mismatch is deleterious, the fitness landscape is assumed to be saddle-like (see Arnegard et al. 2014), where hybrids with mismatched phenotypes are displaced along the steep sides of the saddle and have lower fitness than individuals with relatively ‘matched’ trait values (whether parental or somewhat intermediate). We first consider hybridization between sympatric benthic and limnetic populations of stickleback. These populations, which are reproductively isolated species due to strong assortative mating (Rundle et al. 2000) and reduced hybrid fitness due to extrinsic selection pressures (Hatfield and Schluter 1999), have evolved independently in at least five watersheds in British Columbia, Canada (McKinnon and Rundle 2002; McPhail 1992). Although reproductively isolated in the wild, the species pairs have no known intrinsic barriers that reduce fitness in the lab (Hatfield and Schluter 1999). Second, we consider hybridization between allopatric populations of anadromous and solitary freshwater stickleback. As with the benthic-limnetic species pairs, these populations are recently diverged and can readily hybridize. Our results provide compelling support for the hypothesis that environment-specific hybrid incompatibilities, caused by interactions between individual hybrids and complex ecological environments, exist between recently diverged stickleback populations.

## Methods

### Data sources

We used both previously published and unpublished data in our analyses. Summary information about each data source is listed in Table 1. We base our main inference on a comparison of ancestry heterozygosity in hybrids born and raised in aquaria to hybrids from the same cross types born and raised in experimental ponds, which are semi-natural ecosystems. See Arnegard et al. (2014) and Schluter et al. (2021) for additional information about ponds. Pond-raised crosses capture both ‘intrinsic’ and ‘extrinsic’ incompatibilities, whereas aquarium-raised crosses are expected to capture ‘intrinsic’ incompatibilities that impact hybrid fit-ness.

**Table 1.** Summary of data sources.

| species | design | study | population | method | gen | env. | *n* fish | *n* loci ± [1SD] |
| --- | --- | --- | --- | --- | --- | --- | --- | --- |
| ben × lim | biparental | present | Paxton | microsatellites | F<sub>2</sub> | lab | 89 | 97.1 ± 5.8 |
| ben × lim | biparental | Conte et al. (2015) | Paxton | SNP array | F<sub>2</sub> | pond | 636 | 62.0 ± 0.0 |
| ben × lim | biparental | present | Priest | microsatellites | F<sub>2</sub> | lab | 90 | 22.9 ± 1.1 |
| ben × lim | biparental | Conte et al. (2015) | Priest | SNP array | F<sub>2</sub> | pond | 412 | 89.0 ± 0.0 |
| ben × lim | 8 × F<sub>0</sub> | Arnegard et al. (2014) | Paxton | SNP array | F<sub>2</sub> | pond | 615 | 62.5 ± 17.9 |
| ben × lim | see methods | Bay et al. (2017) | Paxton | SNP array | F<sub>2</sub> | pond | 302 | 183.2 ± 19.7 |
| ben × lim | 4 × biparental | Rennison et al. (2019) | Paxton | GBS (RAD) | F<sub>2</sub> & F<sub>3</sub> | pond | 649 | 85.1 ± 34.0 |
| marine × fresh | biparental | Schluter et al. (2021) | LCR × Cranby | SNP array | F<sub>2</sub> & F<sub>3</sub> | pond | 723 | 120.3 ± 5.6 |
| marine × fresh | biparental | Rogers et al. (2012) | LCR × Cranby | microsatellites | F<sub>2</sub> | lab | 374 | 59.2 ± 4.3 |

Studies raising fish from the same cross type (benthic × limnetic or marine × freshwater) in the same environment (aquaria or pond) were combined for analysis. Two studies (Rennison et al., 2019; Schluter et al., 2021) genotyped both F_2_ and F_3_ hybrids, which we analyze together because we found that this choice does not affect our conclusions (Fig. S2). Similarly, data were analyzed together for four studies of benthic × limnetic hybrids from Paxton Lake raised in experimental ponds because excess ancestry heterozygosity was statistically indistinguishable among them (Fig. S3). This grouping of studies was done only to simplify the presentation of results—patterns are highly repeatable across replicates and analyses showing results for each pond and/or study separately are shown in Fig. S4. Relevant details of each data source are outlined below, but see the original studies for full details including animal use permits.

Studies genotyped fish using either microsatellites, single nucleotide polymorphism (SNP) arrays, or Genotyping-by-Sequencing (GBS). All three lab studies used microsatellites, whereas the pond studies all used SNPs or GBS. We examined potential concerns resulting from different genotyping methods and found no evidence that our results are caused by such differences. First, we only consider loci where parents have no alleles in common and thus can accurately polarize ancestry. Second, we only use loci that were heterozygous for ancestry in F_1_s, so loci with ‘null’ microsatellites or any difficulties in distinguishing alleles would be filtered out (see section on data filtering below). In the largest microsatellite dataset (Rogers et al., 2012), 100 % of loci that were different in parents were accurately called as heterozygous across eight F_1_s (288 of 288 loci across all eight F_1_ fish). SNP genotypes of the same cross type similarly had 100 % heterozygosity in F_1_s (Schluter et al., 2021). Finally, in 1,000 simulations re-sampling our dataset to only a single marker per chromosome, 99.4 % of estimates of our statistical main effect were in the same direction as detected in the full dataset. In light of the above, we suggest our the differences detected between lab and pond datasets reflect biology rather than methodology. Allele frequencies and heterozygosity are shown for all genotyped loci (within a given dataset) in Fig. S5.

One additional difference between the pond and aquarium studies is that pond F_2_ hybrids were a result of natural mating among F_1_ hybrids whereas aquarium F_2_s were produced via artificial crosses. We do not anticipate that this will affect our conclusions, however, because we only consider loci that were fixed differences between F_0_s, and thus are expected to segregate in a 1:2:1 pattern regardless of the process that united eggs and sperm.

#### Benthic × limnetic crosses

We obtained data from four sources for the pond-raised benthic × limnetic hybrids. The data from aquaria are unpublished. Relevant details of each data source are given below. Final sample sizes from each data source are given in Table 1.

Conte et al. (2015) generated a single F_1_ family from each of the Priest and Paxton Lake species pairs. Both were founded by a single wild benthic female and a limnetic male that were collected and crossed in 2009. 35 adult F_1_ Paxton Lake hybrids and 25 adult F_1_ Priest Lake hybrids were released into separate ponds where they bred naturally to produce F_2_ hybrids. F_2_ adults were collected over one year later and were genotyped using a SNP array (Jones et al., 2012). 246 SNPs were found in the Paxton cross and 318 were found in the Priest cross.

Arnegard et al. (2014) conducted a pond experiment with eight F_0_ grandparents from Paxton Lake. Two crosses were between limnetic females and benthic males, and two crosses were between benthic females and limnetic males. Five F_1_ males and five F_1_ females from each family were added to a single pond in March 2008 where they bred naturally. Juvenile F_2_s were collected in October of that same year and genotyped at 408 SNPs using the SNP array.

Bay et al. (2017) genotyped F_2_ hybrid females between Paxton Lake benthics and limnetics. Fish are from several crosses and study designs. One used a cross with four unique F_0s_ that were used to produce two F_1_ families—one with a limnetic as dam and the other with a benthic as dam. A second had eight unique F_0_s, where two F_1_ crosses were in each direction. These two crosses used wild fish collected in 2007. A third set of crosses was done in 2009, one in each direction, then the two F_1_ families were released into separate ponds. Since the goal of the authors’ study was to examine the genetics of mate choice, a large number of F_2_ females were genotyped at a small number of microsatellite markers. A subset of F_2_ females identified in this parentage analysis were genotyped at 494 SNP markers using the SNP array. A total of 302 females were assigned to families with 10 or more full-sibs (which was necessary for linkage mapping).

Finally, Rennison et al. (2019) conducted a study with four unique Paxton Lake benthic × limnetic F_1_ hybrid families that were each split between two ponds. One pond in each pair contained a cutthroat trout predator (heterozygosity did not differ across pond types and data are pooled across all pairs and pond types). Wild fish were caught in 2011 and F_1_s were released in 2012. Fish bred naturally and juvenile F_2_s were sampled in September of that same year. F_3_ hybrids were collected in September 2013. Approximately 50 fish from each pond and hybrid generation were genotyped at over 70,000 loci using restriction-site associated DNA sequencing (genotyping-by-sequencing), and after filtering and selection of diagnostic loci we retained 2243 SNPs.

The Paxton and Priest Lake laboratory cross data are original to this study. Crosses used a single wild-caught benthic female fish and a single wild-caught limnetic male fish as F_0_ progenitors. Wild fish were crossed in 2003. Sibling mating of F_1_ hybrids was used to produce a single F_2_ hybrid family for analysis, and fish were raised in glass aquaria and fed *ad libitum*. 92 Priest Lake F_2_s were genotyped at 85 microsatellite markers, and 86 Paxton Lake F_2_s were genotyped at 216 microsatellite markers. See the supplementary methods for additional details.

#### Marine × freshwater crosses

Schluter et al. (2021) conducted a pond experiment with anadromous (hereafter ‘marine’) × freshwater hybrids. This study crossed a marine female from the Little Campbell River, BC, with a freshwater male from Cranby Lake, BC. Over 600 juvenile F_2_ hybrids were introduced into the ponds directly in August 2006. F_2_s were produced from six F_1_ families—six unique females were crossed with four males (two males were crossed twice each). F_2_s overwintered with an estimated over-winter survival rate of approximately 86 % (from mark-recapture). In spring 2007, surviving F_2_s bred and were genotyped at 1,294 bi-allelic SNP markers using a SNP array. 500 of their F_3_ hybrid offspring were collected in October 2007 and were genotyped with the same methodology.

The data for the laboratory marine × freshwater cross were originally published by Rogers et al. (2012). The population used a single Little Campbell River female as the F_0_ dam and a single Cranby Lake male as the F_0_ sire. Wild adult fish were captured in 2001 to generate a single F_1_ hybrid family. Four F_2_ crosses were made from individuals from this F_1_ family (eight F_1_ parents total). F_2_ hybrid fish were genotyped at 96 microsatellite markers.

### Marker filtering and estimating excess ancestry heterozygosity

For each dataset, we restricted our analysis to loci where the F_0_ progenitors of a given F_2_ family had no alleles in common (e.g., all ‘BB’ in benthic F_0_s and all ‘LL’ in limnetic F_0_s) and where all F_1_ hybrids were heterozygous for ancestry (e.g., all ‘BL’). GBS data were filtered to include SNPs with > 20× coverage for a given individual. Final sample sizes of fish and markers are given in Table 1. In all cases, the sex chromosome (chromosome 19) was not analyzed.

Because some studies have more individuals than loci, and others have more loci than individuals, we analyze ancestry heterozygosity both in individuals (averaged across loci) and at loci (averaged across individuals). We retained individuals for which at least 20 loci were genotyped, and retained loci for which at least 20 individuals were genotyped. Differences in genotyping success and/or family structure caused the number of genotyped loci to differ among individuals for a given study (Table 1). For simplicity, we focus on the analysis of individuals in the main text.

Deviation from the expected 50:50 ancestry proportions in F_2_ hybrids—via variance introduced by the recombination process (or directional selection against one ancestry)— reduce the expected heterozygosity below 0.5. To account for this, we base our main inference on estimates of *excess* ancestry heterozygosity. Excess ancestry heterozygosity was calculated as observed ancestry heterozygosity (*p*_AB_) minus expected ancestry heterozygosity (2*p*_A_p_B_, where *p*_A_ and *p*_B_ are the frequencies of both ancestries at the locus or in the individual’s genome). Our conclusions are unchanged, however, if an uncorrected ‘observed’ heterozygosity is used as the response variable or if the expected heterozygosity is adjusted for sample size (i.e., multiplying by 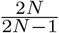 following Irwin et al. 2018; both analyses included in archived R script).

### Data analysis

We evaluated excess ancestry heterozygosity in aquarium and pond studies using one-sample *t*-tests and compared them directly using linear models. Linear models included individual excess ancestry heterozygosity as the response variable and environment (lab or pond) as a categorical predictor. For the benthic × limnetic data, ‘lake’ was included as a second categorical predictor. A lake × environment interaction term was non-significant and was omitted from the final model. Conclusions are unchanged if we analyze excess ancestry heterozygosity of loci averaged across individuals (Fig. S6).

Another possible cause of excess ancestry heterozygosity is genotyping error. Simulations of genotyping error— where all errors are assumed to have resulted in true homozygotes being called as heterozygotes—indicate that error rates in excess of 5% are necessary to cause the pattern we observe (not shown; see archived R script). Given we detected 0 % false homozygosity calls in F_1_ hybrids, we believe our error rate is much smaller than 5%.

## Results

Mean individual excess ancestry heterozygosity was not significantly different from zero in any aquarium-raised stickleback hybrid cross (see left side [red] of panels in Fig. 2). Specifically, one-sample *t*-tests evaluating whether excess ancestry heterozygosity was significantly different from zero failed to reject the null hypothesis for both benthic × limnetic crosses (mean & 95 % CI Paxton Lake—*μ*= 0.014 [−0.010, 0.037], *t*_88_ = 1.16, *P* = 0.25; Priest Lake—*μ* = 0.0098 [−0.014, 0.034], *t*_89_ = 0.81, *P* = 0.42). The aquarium-raised marine × freshwater hybrid cross was similar (*μ* = −0.0047 [−0.013, 0.00396], *t*_373_ = 0.28, *P* = 0.28). Mean individual excess ancestry heterozygosity exceeded zero in each pond-raised cross (see right side [blue] of panels in Fig. 2). This was the case for both Paxton and Priest Lake hybrids for the benthic × limnetic crosses (mean & 95 % CI Paxton Lake—*μ* = 0.029 [−0.024, 0.033], *t*_2222_ = 13.16, *P* < 0.0001; Priest Lake—*μ* = 0.039 [0.029, 0.048], *t*_411_, *P* < 0.0001), as well as the marine × freshwater cross (*μ* = 0.034 [0.027, 0.041], *t*_722_ = 9.33, *P* < 0.0001). In linear models, we found that individual mean excess ancestry heterozygosity was 2.2 % higher in pond-raised benthic limnetic hybrids than in aquarium-raised hybrids (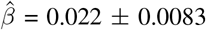 [SE; units are heterozygote frequency], F_1,2509_ = 6.89, *P* = 0.0087) (Fig. 2A; also see Fig S7 for plots of individual hybrid index and heterozygosity). For the marine × freshwater crosses, we found that individual mean excess ancestry heterozygosity was 3.9 % higher in pond-raised fish than in aquarium-raised fish 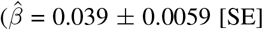, F_1,1095_ = 41.88, *P* = 1.46 × 10*−*10) (Fig. 2B). The signal of excess ancestry heterozygosity was variable among chromosomes, though the majority had values exceeding 0 (Fig. S8).

**Fig. 2.**
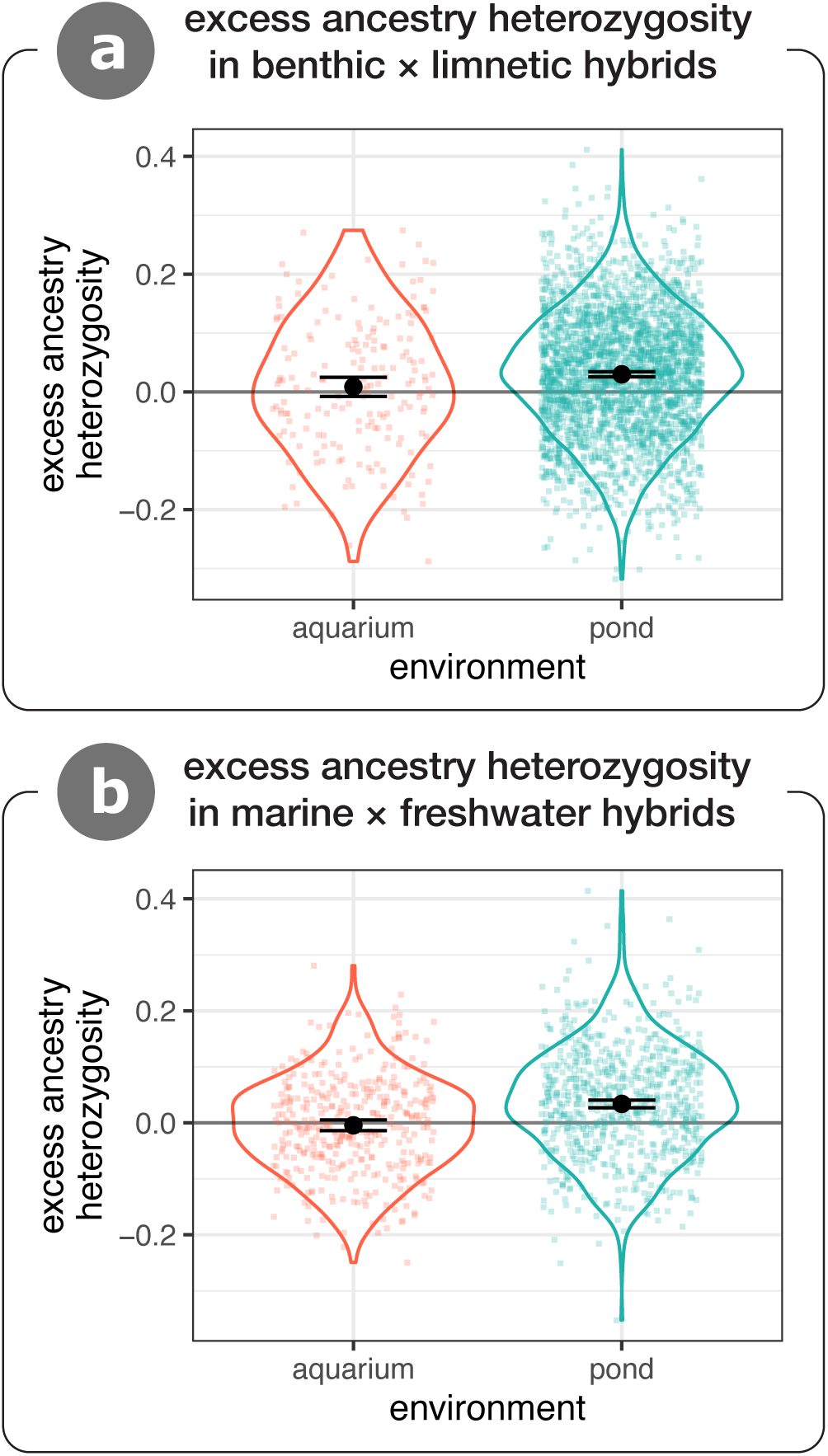
Excess ancestry heterozygosity in recombinant threespine stickleback hybrids in the lab (aquarium) and field (pond). Benthic × limnetic crosses (A) and marine × freshwater crosses (B) are shown separately. Violin overlays show the full distribution of the data and small coloured points show values for individual fish. Large black points are the group means ± 95 % CI.

## Discussion

Ecological selection acting on hybrids is a critical determinant of gene flow between diverging lineages (Schluter, 2000). Yet, detecting how divergent alleles interact to mediate hybrid fitness in relevant ecological contexts has proven difficult due to the small individual effects of interacting loci. Here, we tested whether a coarse-grained signal of selection against hybrid incompatibilities—elevated excess ancestry heterozygosity (Simon et al., 2018)—differed between laboratory and field replicates of crosses between the same populations. We find that excess ancestry heterozygosity was elevated in recombinant stickleback hybrids raised in experimental ponds compared to those from similar crosses raised in aquaria. This result is consistent with the hypothesis that certain ecologically mediated hybrid incompatibilities between recently diverged stickleback populations act more strongly in field settings than in the lab. Our finding implies that individual stickleback with a greater mismatch in parental traits are less likely to survive than those with lesser mismatch. Below, we consider whether other processes could plausibly explain this result, discuss the relevance of our findings for speciation, and highlight opportunities for future research.

### Alternative causes of excess ancestry heterozygosity

The lack of excess ancestry heterozygosity in aquaria is not surprising given what is known about ‘intrinsic’ hybrid incompatibilities in stickleback. Previous studies of benthic × limnetic hybrids have found no evidence for intrinsic inviability in F_2_ benthic × limnetic stickleback crosses using measures of embryo development and hatching success (Hatfield and Schluter, 1999; McPhail, 1984, 1992) or lifetime fitness (Hatfield and Schluter, 1999). In a recent review summarising the literature on reproductive isolation in threespine stickleback, Lackey and Boughman (2017) report that ‘intrinsic’ barriers are typically weak to nonexistent. Both marine (i.e., anadromous) × freshwater and benthic × limnetic crosses had no evidence for intrinsic inviability whereas the authors found evidence for hybrid ecological inviability in both systems (Lackey and Boughman, 2017). Thus, it would have been surprising if we had recovered any signal of selection on excess ancestry heterozygosity in lab-raised hybrids.

Although we hypothesise that selection against trait mismatch (i.e., ecological hybrid incompatibility) causes the observed patterns of selection on ancestry heterozygosity among surviving individuals in the ponds, several other mechanisms could possibly underlie this pattern. These alternative mechanisms—environment-specific heterosis and environment-specific dominance—involve processes operating at single loci rather than interactions among loci. Ultimately, the data presented here have limited ability to conclusively distinguish between single-locus processes like heterosis and multi-locus processes like incompatibilities, but we discuss the strength of evidence for different possible alternative mechanisms below.

Heterosis refers to a case where, for a given locus, the heterozygote has greater fitness than the parental genotypes. This could obviously lead to selection driving excess ancestry heterozygosity. In the benthic × limnetic crosses, environment-specific heterosis can be disregarded based on prior knowledge about hybrid fitness in this system. If heterosis were common, then F_1_ hybrids should have higher fitness than parents. However, F_1_ and reciprocal backcross hybrids have *lower* growth and/or survival than both parent taxa in field experiments (Hatfield and Schluter 1999; Rundle 2002; Vamosi et al. 2000). These patterns are opposite to what would be expected as a result of field-specific heterosis, suggesting that it is unlikely to be acting in the benthic × limnetic crosses. Less is known about heterosis in marine × freshwater crosses. It is conceivable that heterosis could act in the lab via body condition and general health, but does not result in mortality and thus would not affect mean excess ancestry heterozygosity. However, the growth rate of F_1_ benthic × limnetic hybrids in the lab matches the additive expectation of parents (Hatfield, 1997; Hatfield and Schluter, 1999). In the present study, we find no relationship between body size and excess ancestry heterozygosity in any of the aquarium raised crosses (Fig. S9; family sizes are too small for a robust analysis in ponds; see Arnegard et al. (2014) for discussion of using body size as a component of fitness). Thus, we conclude that pond-specific heterosis is unlikely to explain higher ancestry heterozygosity in lab versus pond raised stickleback hybrids.

We can also use predictions that are specific to heterosis or hybrid incompatibilities to differentiate between them. Specifically, heterosis depends on interactions within a locus while incompatibilities which depend on interactions between loci. If the benefit of ancestry heterozygosity was due to interactions within loci alone, we would expect to see no relationship between genome wide admixture proportion (i.e., hybrid index) and excess heterozygosity—all heterozygosity is useful. By contrast, if excess heterozygosity was attributable to interactions among loci as in the hybrid incompatibility model we would expect diminishing benefits of excess ancestry heterozygosity as genome-wide ancestry proportions became more parent-like (i.e., as hybrid index deviates from 0.5). As expected by the incompatibility model, excess heterozygosity declines as the hybrid index of pond-raised individuals deviates from 0.5 (Spearman’s *ρ* = −0.059; *P* = 0.0006), while there is no relationship in the lab (*ρ* = −0.012; *P* = 0.77) (Fig. S10). While this observation does not eliminate the possibility that some of the observed excess heterozygosity is driven by environment-specific heterosis, it is consistent with our hypothesis that excess heterozygosity largely results from selection against extrinsic incompatibilities.

Finally, an additional analysis indirectly provides evidence of hybrid incompatibilities in the data from Arnegard et al. (2014). Specifically, Arnegard et al. (2014) classified F_2_ hybrids into four groups (‘A’, ‘B’, ‘L’, and ‘O’) based on individual niche use. ‘B’, ‘O’, and ‘L’ individuals had benthic-like, intermediate, and limnetic-like diets, respectively. ‘A’ individuals, however, had unusual diets, were smaller, and had a greater extent of mismatched trait combinations compared to the other groups. The authors hypothesise that trait mismatch caused these fish to grow more slowly than more ‘matched’ individuals. Our analysis reveals that ‘A’ group individuals have lower excess ancestry heterozygosity than non-‘A’ individuals (Fig. S11)—as expected if lower excess ancestry heterozygosity correlates with higher trait mismatch. This reanalysis suggests a link between trait mismatch, ancestry heterozygosity, and fitness in stickleback hybrids raised in natural environments.

### Relation to other studies of incompatibilities and ancestry heterozygosity

Our results contribute to a growing understanding of the biology of environment-dependent hybrid incompatibilities. In natural hybrid populations of swordtail fishes, Schumer et al. (2014) estimate that dozens of incompatibilities separate parent species (Schumer and Brandvain, 2016). The authors also suggest that many of these are likely subject to natural or sexual selection (Schumer et al., 2014). Previous studies on hybrid stickleback (Arnegard et al., 2014; Keagy et al., 2016) have estimated fitness landscapes that are consistent with the hypothesis that mismatched trait combinations are selected against, and our analysis of genetic data supports this hypothesis. More broadly, our results are consistent with predictions generated from theoretical models of speciation and adaptation (e.g., Fisher’s [1930] geometric model; Simon et al. 2018).

Our findings also highlight differences from previous analyses of selection on hybrids in different environments. In yeast, selection for *low* ancestry heterozygosity is common in hybrids when tested in the lab (Lancaster et al., 2019; Zhang et al., 2020). This difference might result from the fact the lab media that yeast were raised in are novel environments, and transgressive traits suited to these environments result from excess ancestry homozygosity (see Fig. 1 and Fig. S1). Stickleback populations in post-glacial lakes are specialists on zooplankton or benthic invertebrates when they coexist with a competing fish species (Miller et al., 2019; Schluter, 1995; Schluter and McPhail, 1992), or are generalist populations that make use of both niches (Bolnick and Ballare, 2020). In contrast to other systems like Caribbean pupfish where scale-eating and mollusc-feeding have evolved (Martin and Wainwright, 2013), no novel niches are colonised by stickleback in post-glacial lakes and therefore transgressive phenotypes are expected to be deleterious. Qualitative patterns of selection against hybrids driving excess ancestry heterozygosity might therefore depend on the availability and nature of novel ecological niches.

### Outlook, caveats, and conclusions

While we identify genetic signatures consistent with the existence of environment-specific hybrid incompatibilities, we cannot begin to identify their specific mechanisms, including when they arise during ontogeny, without connecting phenotype to genotype. Experiments that directly manipulate individual phenotypes, or interactions between individuals and their environments, are needed to establish such causality. As a field, we should aim to identify the types of traits that typically underlie ecological hybrid incompatibilities. Integrating field studies of hybrid incompatibility with QTL mapping of ecologically important traits (Arnegard et al., 2014; Schemske and Bradshaw, 2002) represents an exciting new frontier for empirical research into the mechanisms of speciation. Moreover, our data only allows us to scratch the surface of how incompatibilities are spread across the genome—it will be valuable for future studies with higher resolution genomic data to investigate this further.

We also do not know the strength of selection against ecological hybrid incompatibilities. Simple simulations (included in archived R scripts) illustrate that the strength of selection necessary to generate 3 % excess ancestry heterozygosity in a population of F_2_ hybrids (similar to that observed in the present study) can vary by orders of magnitude depending on assumptions about the genetic architecture of selection. Because the expression of maladaptive trait combinations is predicted to increase with the magnitude of divergence between parent populations (Barton, 2001; Chhina et al., 2021; Thompson, 2020), we may predict that the strength of selection against ecological incompatibilities will increase with the magnitude of divergence between parents. However, quantifying the ecological basis of incompatibilities and their genetic structure will remain technically challenging.

The evidence presented here is consistent with the hypothesis that extrinsic hybrid incompatibilities are an important mechanism of post-zygotic isolation in this system. Our results imply that selection against ecologically-mediated hybrid incompatibilities is active from the earliest stages of divergence. Speciation is largely complete when divergent lineages can stably coexist in sympatry, as is the case for the benthic and limnetic stickleback species pairs. Reproductive isolation between them is primarily thought to have arisen incidentally as a by-product of phenotypic divergence (Conte and Schluter, 2013; Hatfield and Schluter, 1999; Nagel and Schluter, 1998; Rundle, 2002), with additional selection favouring the reinforcement of pre-mating barriers as a result of low hybrid fitness (Hatfield and Schluter, 1999; Rundle and Schluter, 1998). Thus, our results are consistent with the idea that selection against trait mismatch, or ecological hybrid incompatibilities, are associated with extrinsic post-zygotic isolation in this classic system of rapid and recent adaptive radiation.

## Acknowledgements

Feedback from D. Irwin, S. Otto, L. Rieseberg, and R. Stelkens improved the manuscript. Discussion with the Schluter Lab at the University of British Columbia, and Schumer lab at Stanford University improved the analysis. We are grateful to the authors of the primary studies from which we gathered data for their data stewardship. R. Henriques created the bioRxiv LATEXtemplate.

## Author contributions

KAT formulated the idea to compare heterozygosity between lab and pond populations, and the approach taken was refined through discussions with CLP, MDM, DS, and MS. YB contributed analyses to distinguish single- and multi-locus selection. KAT conducted the analysis with input from other authors. AYKA and THV generated the original data from Priest Lake lab-raised hybrids, and AKG and ARW generated the original data from the Paxton Lake lab-raised hybrids. DJR processed and contributed GBS data. KAT wrote the first draft of the manuscript, and all authors contributed to revisions.

## Data accessibility

All data and analysis code used in this article will be deposited in a repository (e.g., Dryad) following acceptance. They will be available to reviewers following submission of this manuscript to a journal.

## Supplementary methods

### Simulations underlying conceptual figures

We used simple simulations in Figure 1 of the main text to illustrate the mechanistic relationship between trait mismatch and heterozygosity. Similar results have been noted elsewhere (Barton, 2001; Simon et al., 2018) but we give our detailed methods herein. We consider the following life history: a single population adapts to a novel environment and then hybridizes with the ancestral population. The phenotype and genotype (hybrid index and heterozygosity) are recorded for F_2_ hybrids.

We use the framework of Fisher’s (1930) geometric model, wherein the phenotype of an organism is a vector of *m* traits, **z** = [*z*_1_*, z*_2_…, *z_m_*]. For simplicity and ease of visualization, we only consider *m* = 2 in the main text (see Fig. S1 for *m* = 10). We assume that mutations are sufficiently rare that they individually sweep through an otherwise monomorphic population. We also assume that mutations all occur at unique loci with free recombination among them (i.e., no linkage) (Kimura, 1965, 1969). Mutations influence the phenotype additively, and are vectors of length *m* where values are drawn from a random normal distribution with a mean of 0 and a standard deviation of *α* (*α* = 0.15). The fitness of a given population is calculated as *w* = exp(−*σ* ‖ **z − o**‖^2^), where ‖**z − o**‖ is the Euclidean distance between the populations current phenotype (**z**) and the optimum (**o**), and sigma is the strength of selection (*σ* = 10). The original phenotype of the population is *z*_0_ = [0, 0] and the optimum phenotype is *z*_opt_ = [1,1]. The selection coefficient *s* value when they arise, where 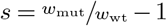. The probability that a given mutation fixes, *π*, is calculated as 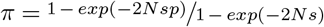, where *N* is the effective population size (*N* = 1000), *p* is the frequency of the mutation in the population 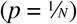, and *s* is the selection coefficient.

We allow 1000 mutations to arise in our simulations, which is sufficient for the adapting population to reach the optimum. After adaptation, we simulate hybridization with the ancestral population (which contains no mutations and has a value of 0 for both traits). We generate 500 F_2_ hybrids, which inherit homozygous ancestral, heterozygous, or homozygous derived ancestry at each locus with probabilities 0.25:0.5:0.25. The genotype of these hybrids determines their phenotype for both traits; these phenotypes are plotted in Fig. 2A. We consider two orthogonal properties of the phenotype: hybrid index and heterozygosity. Hybrid index is the fraction of alleles an individual inherited from the derived parent, and heterozygosity is the fraction of loci that are heterozygous. We finally calculated the Euclidean phenotypic distance from each hybrid’s phenotype to the line connecting parent phenotypes. This distance is the individual hybrid’s ‘mismatch’ as calculated elsewhere (Thompson et al., 2021).

### Genotyping of lab-raised benthic× limnetic hybrids

Ninety-two F2s from the Priest Lake cross were genotyped at 84 microsatellite markers, and 86 F2s from the Paxton Lake cross were genotyped at 216 microsatellite markers following Peichel et al. (2001). We constructed a combined linkage map between the two families using the map integration function in JoinMap 3.0 Van Ooijen and Voorrips 2001. The average centiMorgan (cM) distance between the markers in the Priest Lake and Paxton Lake maps was 15.6 1.85, and 4.5 0.51 (mean 1 SE) respectively. Use of animals was approved by UBC’s Animal Care Committee (A97-0298).

### Supplementary figures

**Fig. S1.**
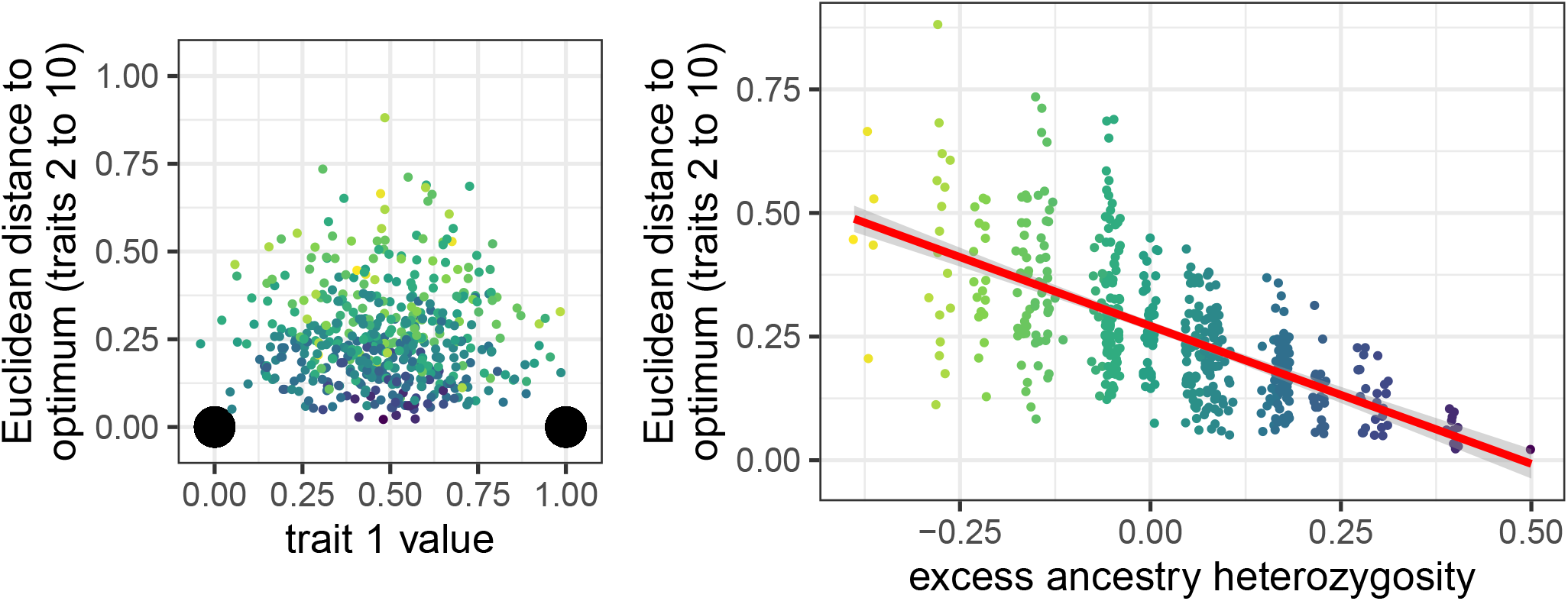
Simulation model illustrating the incompatibility–heterozygosity relationship. The model is as in Fig. 1 in the main text except there are ten traits instead of two and the optimum of the adapting population is ‘0’ for traits 2–9. Plots and model are inspired by Fig. 1 in Barton (2001). Both panels depict results from a representative simulation run of adaptive divergence and hybridization between two populations. Coloured points are individual hybrids, with darker colours indicating higher heterozygosity. Panel (A) depicts the distribution of 500 F_2_ hybrid phenotypes where the *x* -axis depicts the value of the selected trait, and the *y* -axis depicts the Euclidean distance from the optimum for traits 2–9 (i.e., 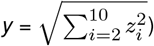. Large black points are the two parent phenotypes. Panel (B) depicts the relationship between excess ancestry heterozygosity and the maladaptive distance from the optimum for individual hybrids (Thompson et al., 2021). Points are slightly jittered horizontally. The plot shows that this maladaptive trait expression is lower in F_2_s with greater excess ancestry heterozygosity. Heterozygosity values are fairly discrete because a small number of loci underlie adaptation in the plotted simulation run.

**Fig. S2.**
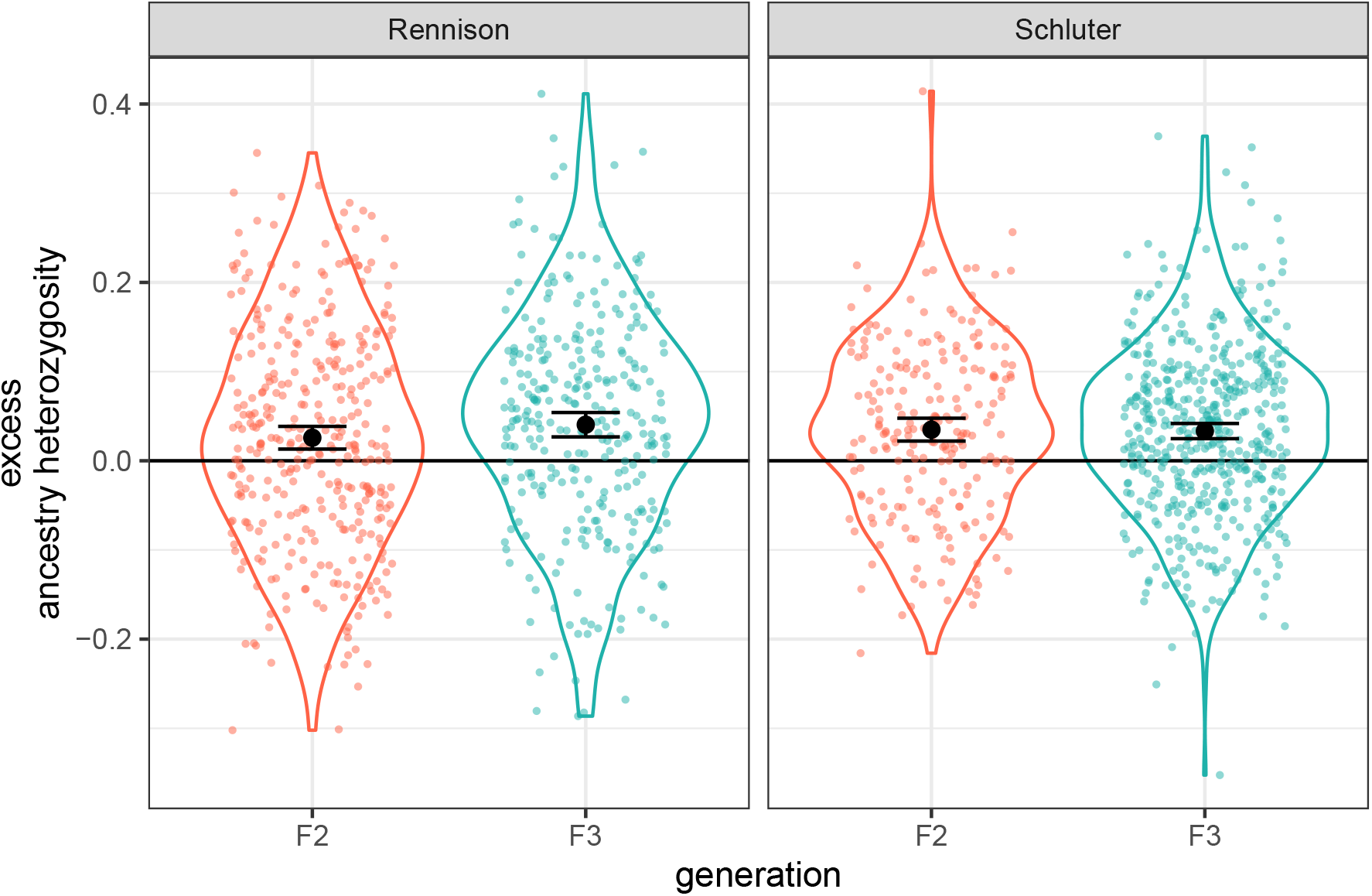
Excess ancestry heterozygosity does not differ between F_2_ and F_3_ hybrids. The plots show individual excess ancestry heterozygosity from the two studies that genotyped both the F_2_ and F_3_ generations (Rennison et al., 2019; Schluter et al., 2021). The means (black dots, ± 95 % CI) do not differ between generations in either study (Rennison—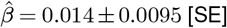, *F*_1,667_ = 2.35, *P* = 0.13; Schluter—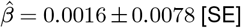, *F*_1,721_ = 0.042, *P* = 0.84).

**Fig. S3.**
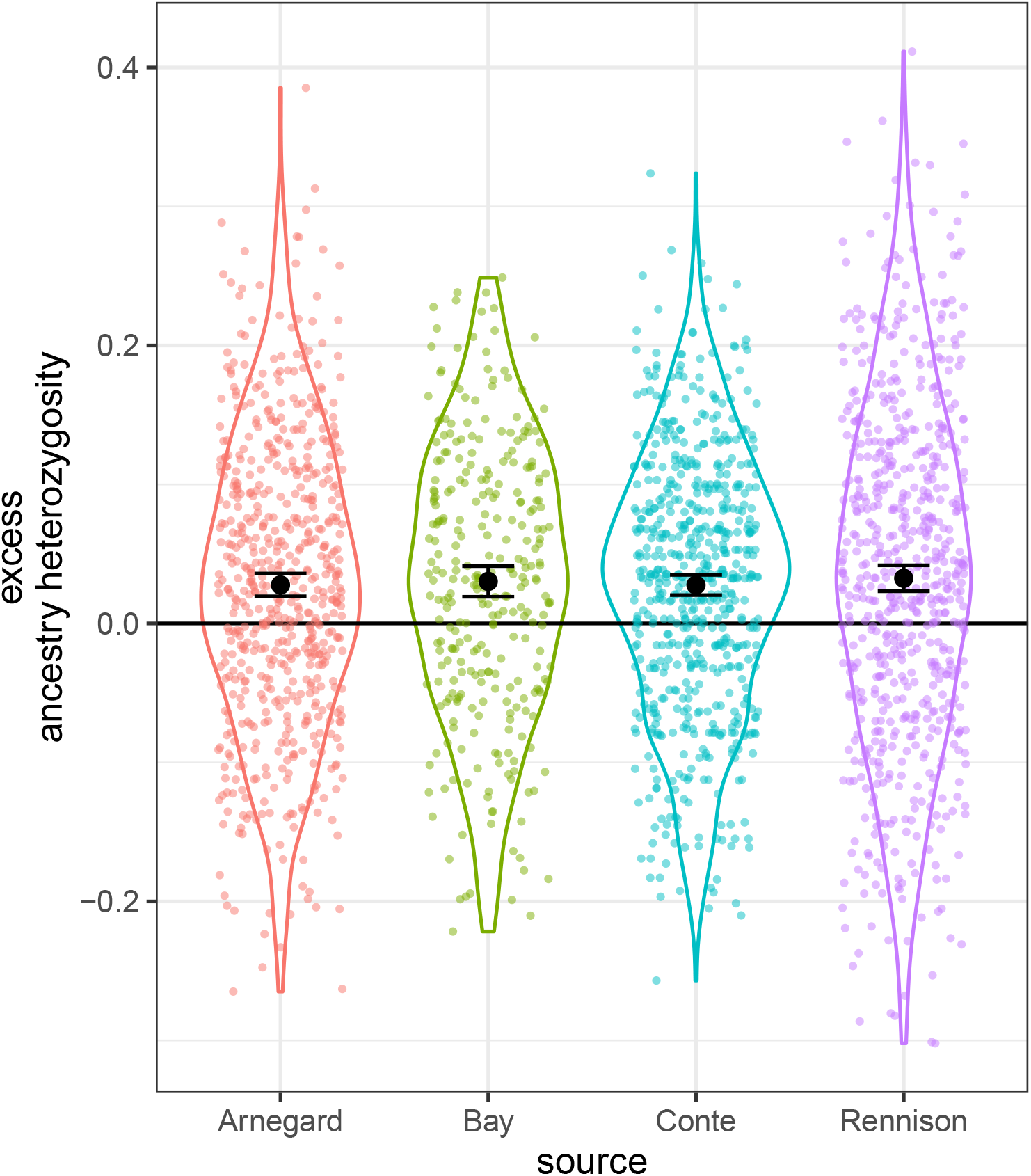
Excess ancestry heterozygosity does not differ between studies involving Paxton Lake benthic × limnetic hybrids. Points are individual hybrids. The means (black dots, ± 95% CI) of all four studies are statistically indistinguishable (*F*_3,2217_ = 0.3036; *P* = 0.82).

**Fig. S4.**
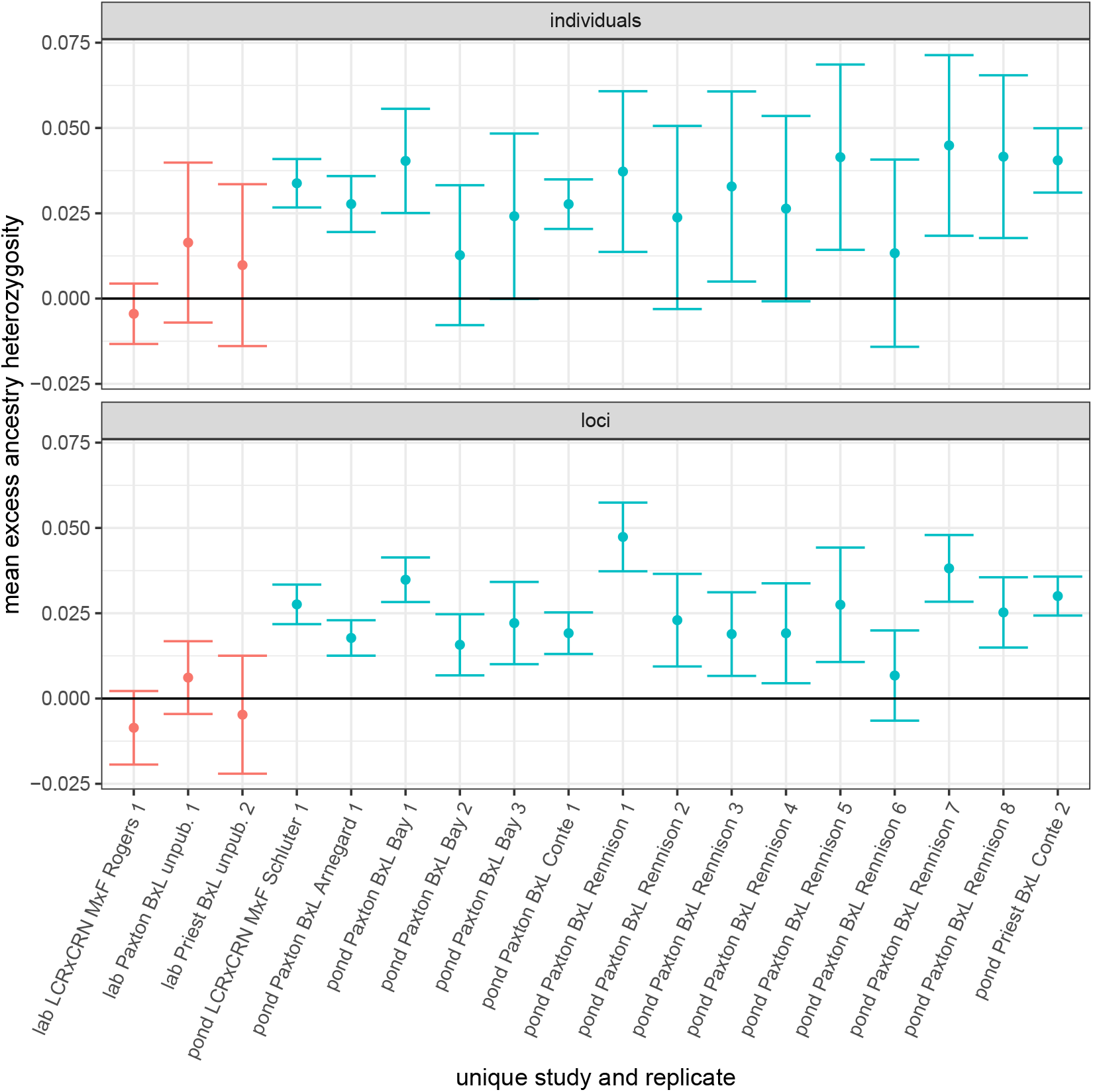
Estimates of mean (± 95% CI) excess ancestry heterozygosity for individuals and loci across ‘replicates’. We consider a replicate to be a unique bi-parental F_0_ cross for aquarium studies, and a unique pond for pond studies. Mean excess ancestry heterozygosity is shown for each such replicate for both individuals (upper) and loci (lower). In each panel the horizontal line indicates no excess ancestry heterozygosity. Red points are ‘lab’ replicates, and blue points are ‘pond’ replicates.

**Fig. S5.**
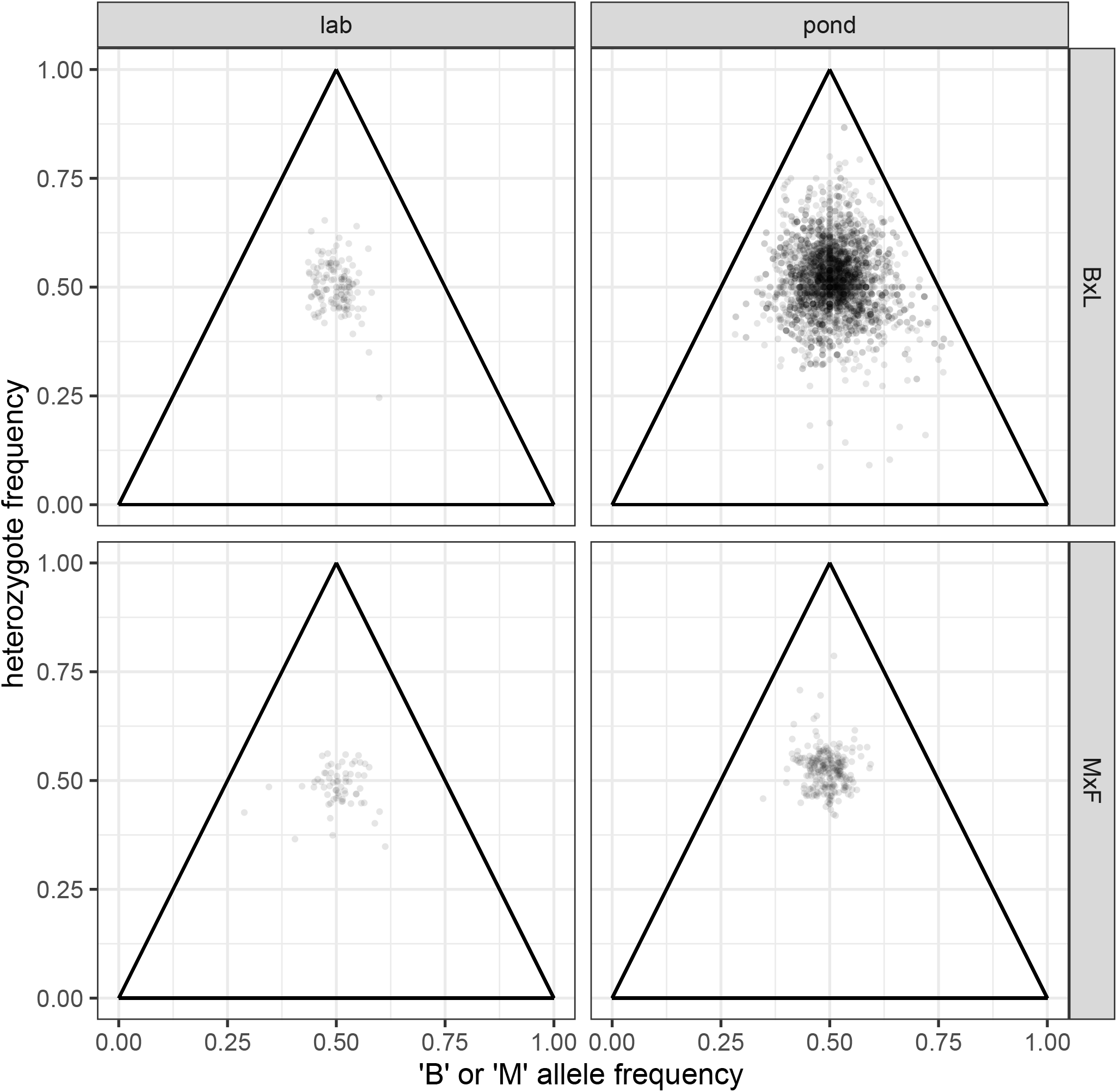
de Finetti ternary diagrams for genotyped *loci*. Each point represents genotyped locus within a given study (i.e., line in Table 1 in the main text) and shows the frequency of either benthic or marine alleles on the *x* -axis and its heterozygosity on the *y* -axis. These graphs are not used for analysis, but rather are shown to allow readers to visualise the structure of the raw data that underlies our analysis.

**Fig. S6.**
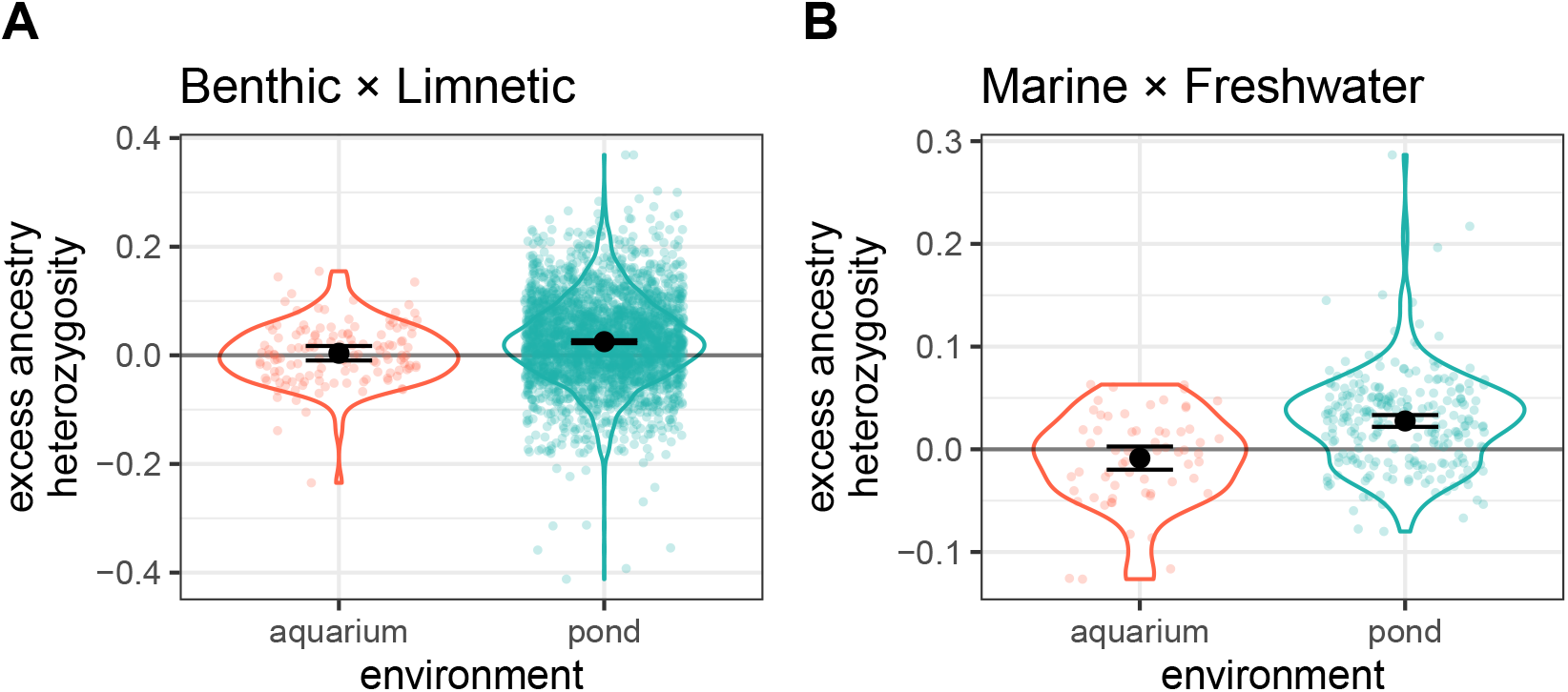
Test of main hypothesis with loci instead of individuals. Each point is a locus genotyped within a given study. Results are qualitatively the same as in the test of individuals. However, this analysis is less robust because loci are not independent in the way that individual fish are. Panel (A) is benthic × limnetic data (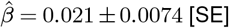, *F*_1,3106_ = 9.50, *P* = 0.0021) and panel (B) is marine × freshwater data (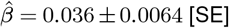, *F*_1,308_ = 31.51, *P* < 0.0001).

**Fig. S7.**
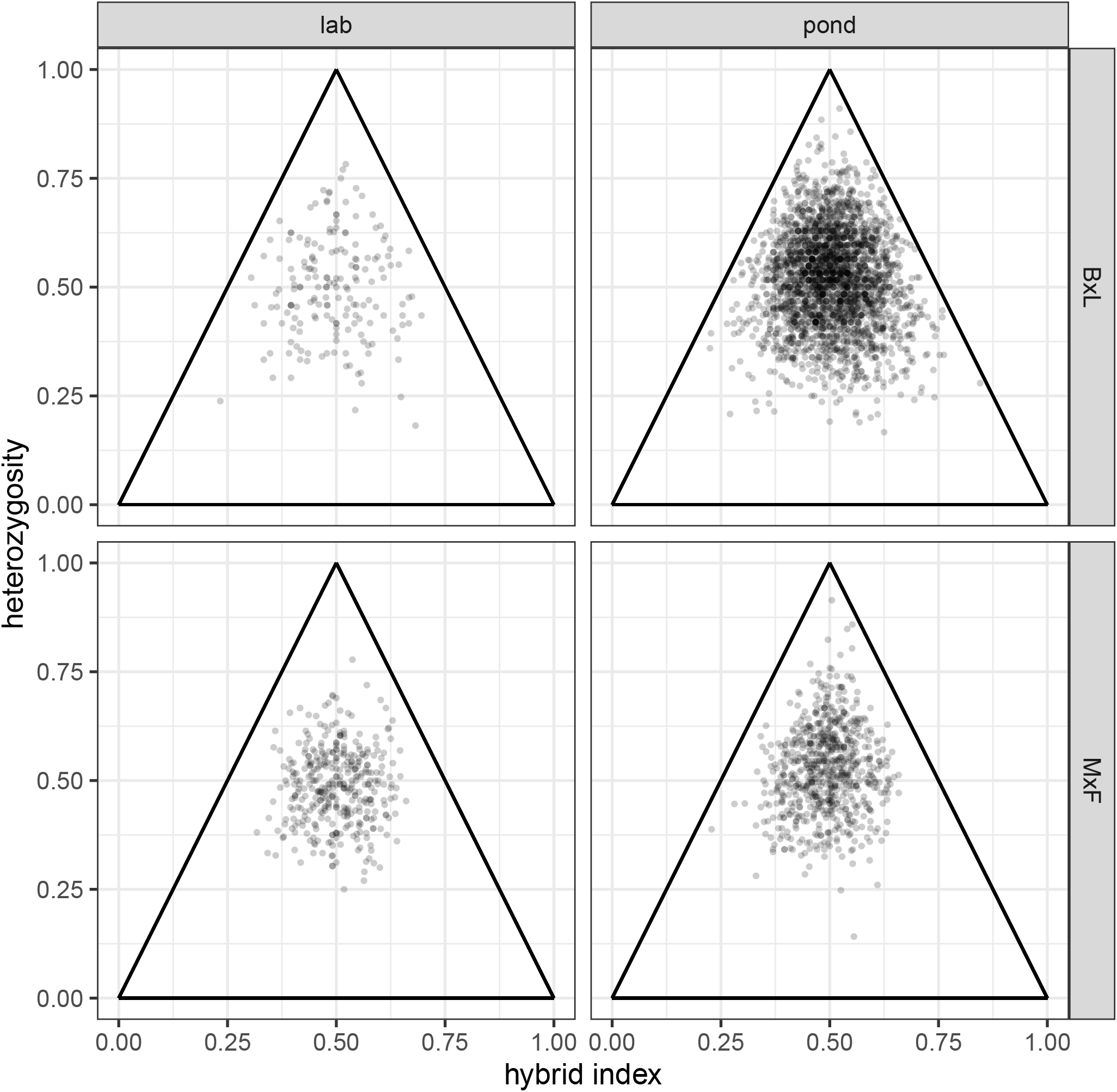
de Finetti ternary diagrams for genotyped *individuals*. Each point represents an individual hybrid and shows each individual’s hybrid index (frequency of benthic or marine alleles in its genome) and its mean heterozygosity. Hybrid index and heterozygosity are used because many loci are being considered simultaneously. These graphs are not used for analysis, but rather are shown to allow readers to visualise the structure of the raw data that underlies our analysis.

**Fig. S8.**
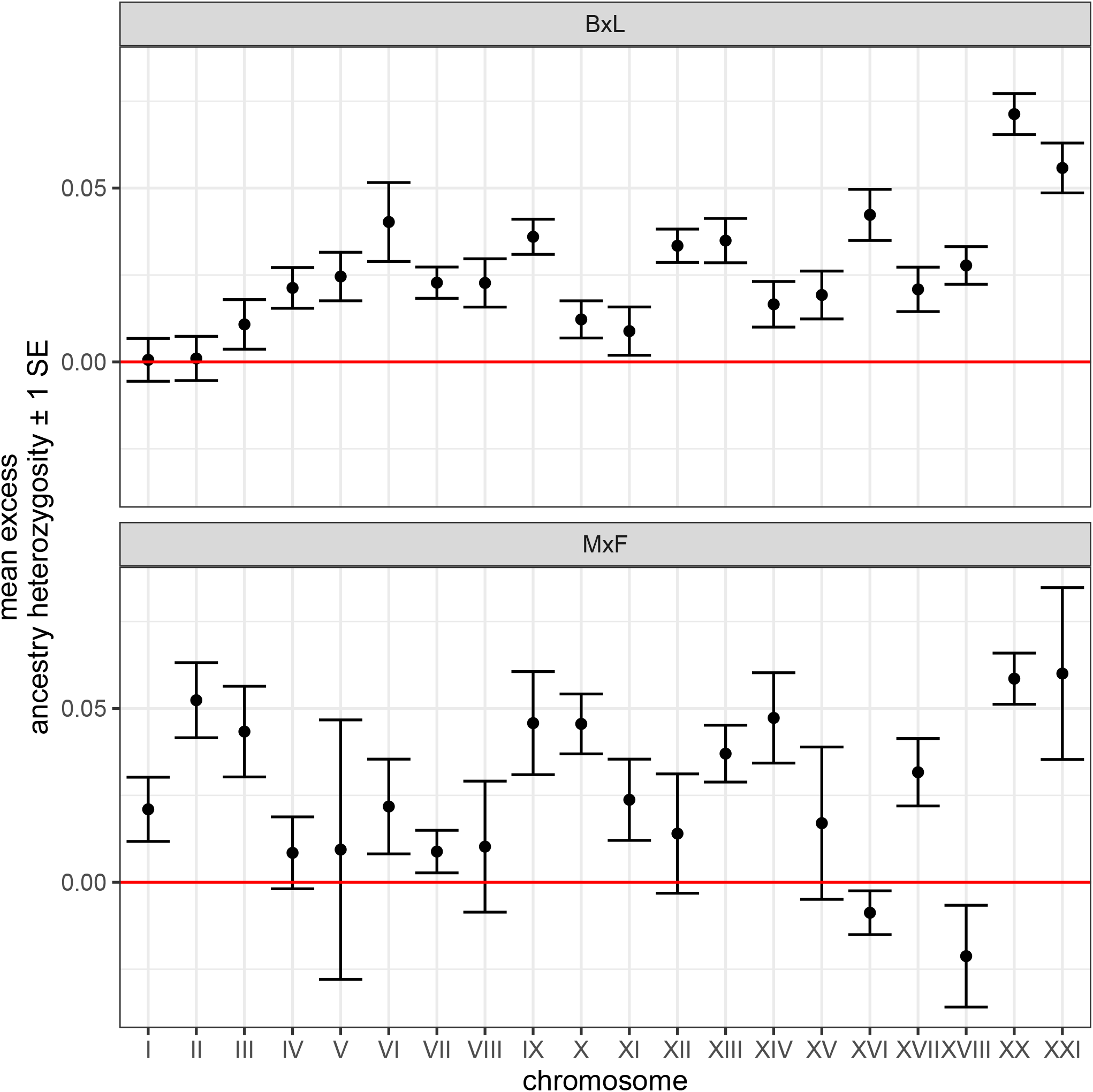
Mean excess ancestry heterozygosity across chromosomes. Each point is the average excess ancestry heterozygosity for all loci on a given chromosome. Linkage group XIX, contains the sex determining region and is not shown. Error bars are 1SE.

**Fig. S9.**
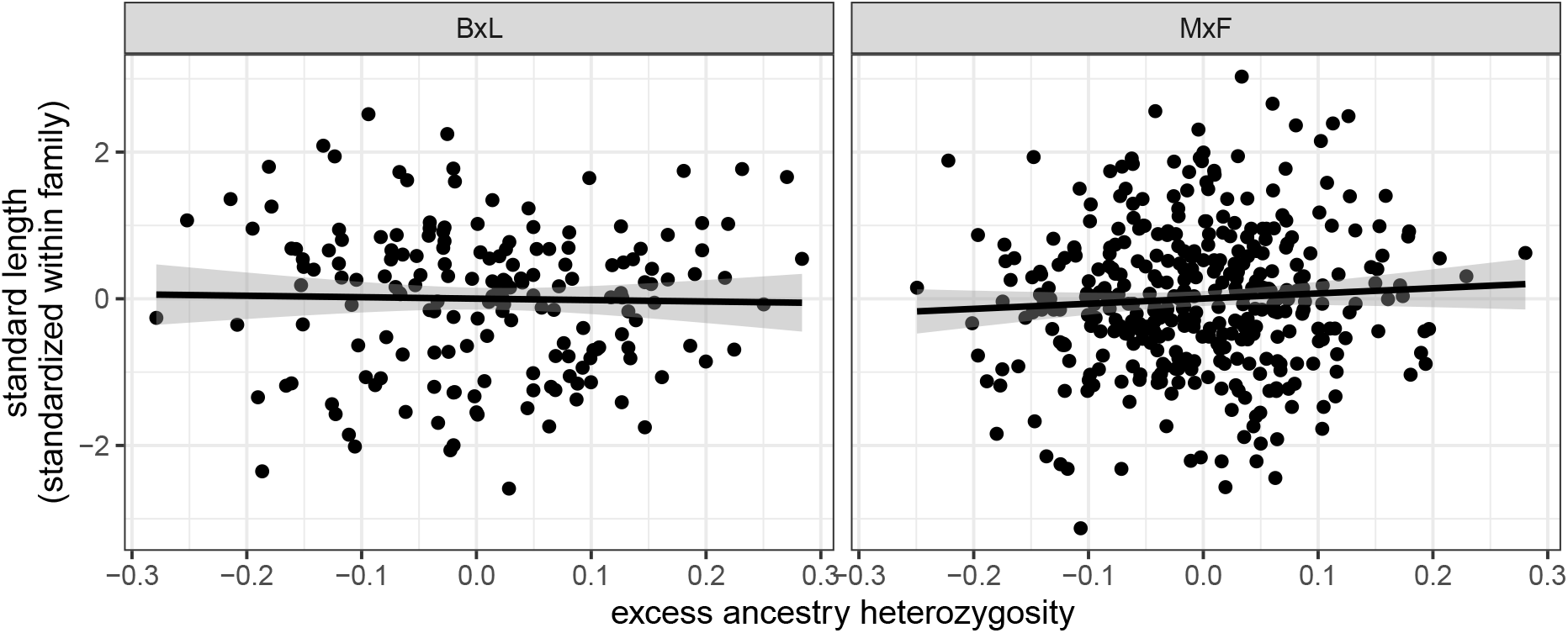
No relationship between individual mean heterozygosity and growth (standard length) in the aquarium-raised bi-parental benthic-limnetic F_2_ hybrids. Results are residuals from `visreg` (Breheny and Burchett, 2017). Each point is an individual F_2_ hybrid. Standard length is standardized within family (one family each for Paxton and Priest lakes for B×L and four families for M×F. The interaction between lake-of-origin mean heterozygosity was non-significant so we plot the main effect across both lakes-of-origin (Paxton and Priest). Mean heterozygosity was not significantly associated with standard length for either cross 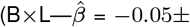 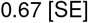, *F*_1,174_ = 0.0075, *P* = 0.93; 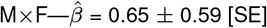, *F*_1,372_ = 1.24, *P* = 0.26). Analyses considering body depth (either individually or in a combined metric of ‘overall size’) give the same qualitative result (not shown).

**Fig. S10.**
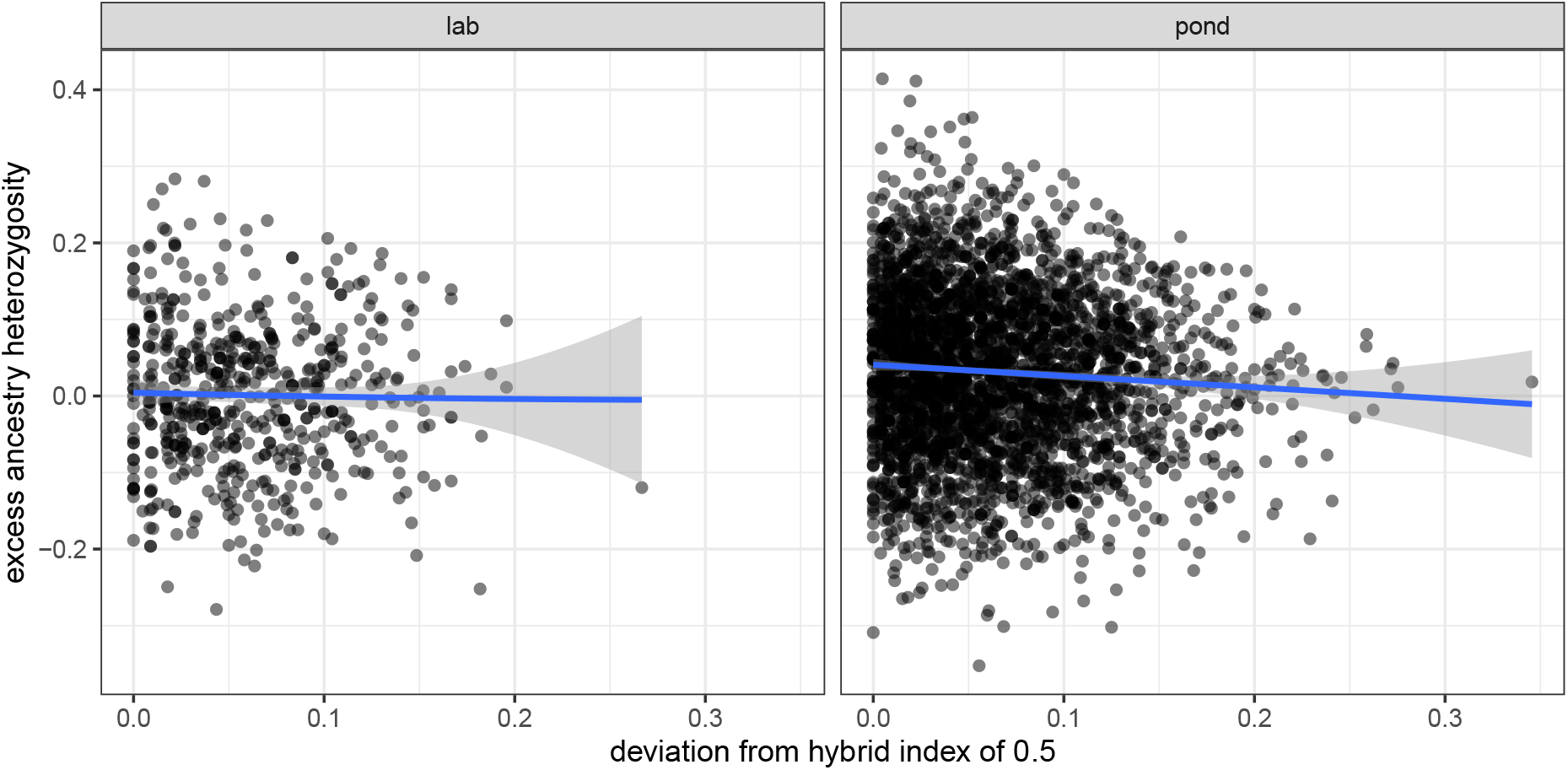
The benefit of excess heterozygosity declines with deviations from a hybrid index of 0.5 in ponds but not in the lab. Each point is an individual recombinant hybrid and data are pooled across cross types. Excess ancestry heterozygosity declines as the hybrid index of pond-raised individuals deviates from 0.5 (Spearman’s *ρ* = −0.059; *P* = 0.0006), while there is no relationship in the lab (*ρ* = −0.012; *P* = 0.77). Bootstrap tests indicate that these two correlations are statistically indistinguishable, so we consider this analysis to be interesting and consistent with our hypothesis, but not conclusive.

**Fig. S11.**
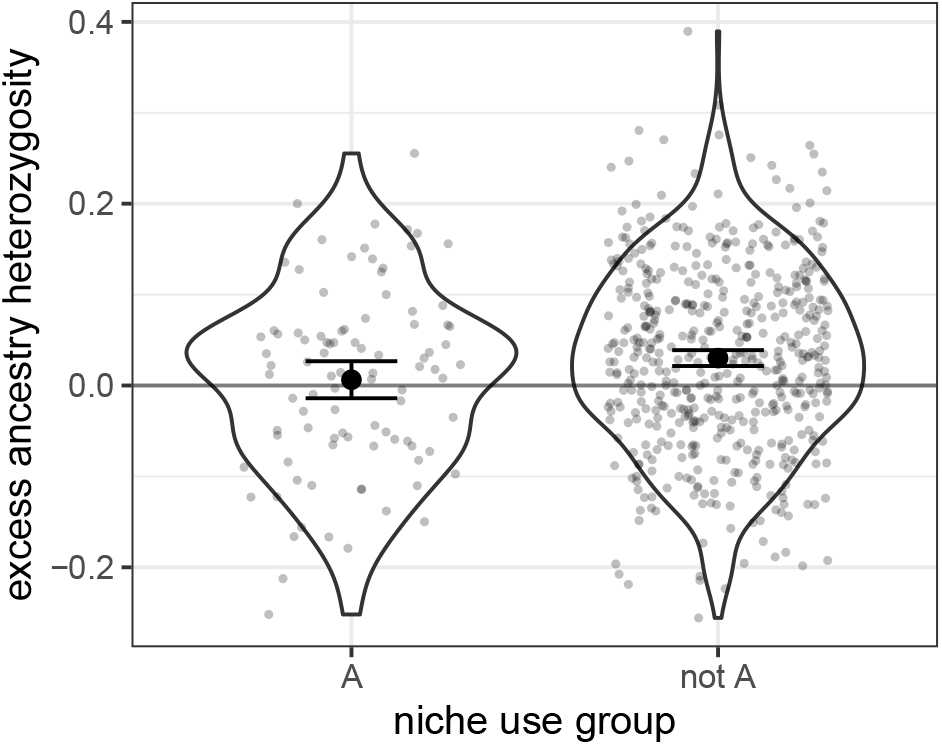
Fish assigned *a priori* as ‘phenotypically mismatched’ (‘A’ group) have lower excess ancestry heterozygosity than non-mismatched fish. Assignments are from Arnegard et al. (2014) and methods are described therein. Each point is an individual F_2_ hybrid. This result implies that phenotypically mismatched individuals have lower excess ancestry heterozygosity than non-mismatched individuals.

## Notes

### Competing Interest Statement

The authors have declared no competing interest.

### Summary of Updates

Fixing silly typo in figure label, and correcting the name of the primary author on bioRxiv website.

## Bibliography

Arnegard, M. E., M. D. McGee, B. Matthews, K. B. Marchinko, G. L. Conte, S. Kabir, N. Bedford, S. Bergek, Y. F. Chan, F. C. Jones, D. M. Kingsley, C. L. Peichel, and D. Schluter. 2014. Genetics of ecological divergence during speciation. Nature 511:307–311.

Barton, N. H. 2001. The role of hybridization in evolution. Molecular Ecology 10:551–568.

Barton, N. H., and K. S. Gale. 1993. Genetic analysis of hybrid zones. Pages 13–45 *in* R. G. Harrison, ed. Hybrid zones and the evolutionary process.

Bay, R. A., M. E. Arnegard, G. L. Conte, J. Best, N. L. Bedford, S. R. McCann, M. E. Dubin, Y. F. Chan, F. C. Jones, D. M. Kingsley, D. Schluter, and C. L. Peichel. 2017. Genetic Coupling of Female Mate Choice with Polygenic Ecological Divergence Facilitates Stickleback Speciation. Current Biology 27:3344–3349.

Bolnick, D. I., and K. M. Ballare. 2020. Resource diversity promotes among-individual diet variation, but not genomic diversity, in lake stickleback. Ecology Letters 23:495–505.

Breheny, P., and W. Burchett. 2017. Visualization of regression models using visreg. R Journal 9:56–71.

Chhina, A. K., K. A. Thompson, and D. Schluter. 2021. Adaptive divergence and the evolution of hybrid trait mismatch in threespine stickleback. bioRxiv.

Conte, G. L., M. E. Arnegard, J. Best, Y. F. Chan, F. C. Jones, D. M. Kingsley, D. Schluter, and C. L. Peichel. 2015. Extent of QTL reuse during repeated phenotypic divergence of sympatric threespine stickleback. Genetics 201:1189–1200.

Conte, G. L., and D. Schluter. 2013. Experimental confirmation that body size determines mate preference via phenotype matching in a stickleback species pair. Evolution 67:1477–1484.

Corbett-Detig, R. B., J. Zhou, A. G. Clark, D. L. Hartl, and J. F. Ayroles. 2013. Genetic incompatibilities are widespread within species. Nature 504:135–137.

Coyne, J. A., and H. A. Orr. 2004. Speciation. Sinauer.

Demuth, J. P., and M. J. Wade. 2007. Population differentiation in the beetle *Tribolium castaneum*. I. Genetic architecture. Evolution 61:494–509.

Fisher, R. A. 1930. The Genetical Theory of Natural Selection. Oxford University Press, Oxford, UK.

Fishman, L., and A. L. Sweigart. 2018. When Two Rights Make a Wrong: The Evolutionary Genetics of Plant Hybrid Incompatibilities. Annual Review of Plant Biology 69:707–731.

Hatfield, T. 1997. Genetic divergence in adaptive characters between sympatric species of stickleback. The American Naturalist 149:1009–1029.

Hatfield, T., and D. Schluter. 1999. Ecological speciation in sticklebacks: environment-dependent hybrid fitness. Evolution 53:866–873.

Irwin, D. E., B. Milá, D. P. Toews, A. Brelsford, H. L. Kenyon, A. N. Porter, C. Grossen, K. E. Delmore, M. Alcaide, and J. H. Irwin. 2018. A comparison of genomic islands of differentiation across three young avian species pairs. Molecular Ecology 27:4839–4855.

Jones, F. C., Y. F. Chan, J. Schmutz, J. Grimwood, S. D. Brady, A. M. Southwick, D. M. Absher, R. M. Myers, T. E. Reimchen, B. E. Deagle, D. Schluter, and D. M. Kingsley. 2012. A genome-wide SNP genotyping array reveals patterns of global and repeated species-pair divergence in sticklebacks. Current Biology 22:83–90.

Keagy, J., L. Lettieri, and J. W. Boughman. 2016. Male competition fitness landscapes predict both forward and reverse speciation. Ecology Letters 19:71–80.

Kimura, M. 1965. A stochastic model concerning the maintenance of genetic variability in quantitative characters. Proceedings of the National Academy of Sciences of the United States of America 54:731–736.

Kimura, M. 1969. The number of heterozygous nucleotide sites maintained in a finite population due to steady flux of mutations. Genetics 61:893–903.

Lackey, A. C., and J. W. Boughman. 2017. Evolution of reproductive isolation in stickleback fish. Evolution 71:357–372.

Lancaster, S. M., C. Payen, C. S. Heil, and M. J. Dunham. 2019. Fitness benefits of loss of heterozygosity in Saccharomyces hybrids. Genome Research 29:1685–1692.

Maheshwari, S., and D. A. Barbash. 2011. The Genetics of Hybrid Incompatibilities. Annual Review of Genetics 45:331–355.

Martin, C. H., and P. C. Wainwright. 2013. Multiple fitness peaks on the adaptive landscape drive adaptive radiation in the wild. Science 339:208–11.

Matute, D. R., I. A. Butler, D. A. Turissini, and J. A. Coyne. 2010. A test of the snowball theory for the rate of evolution of hybrid incompatibilities. Science 329:1518–1521.

McKinnon, J. S., and H. D. Rundle. 2002. Speciation in nature: the threespine stickleback model systems. Trends in Ecology & Evolution 17:480–488.

McPhail, J. D. 1984. Ecology and evolution of sympatric sticklebacks (Gasterosteus): morpho-logical and genetic evidence for a species pair in Enos Lake, British Columbia. Canadian Journal of Zoology 62:1402–1408.

McPhail, J. D. 1992. Ecology and evolution of sympatric sticklebacks (Gasterosteus): evidence for a species-pair in Paxton Lake, Texada Island, British Columbia. Canadian Journal of Zoology 70:361–369.

Miller, S. E., M. Roesti, and D. Schluter. 2019. A Single Interacting Species Leads to Widespread Parallel Evolution of the Stickleback Genome. Current Biology 29:530–537.

Moyle, L. C., and T. Nakazato. 2010. Hybrid incompatibility “snowballs” between Solanum species. Science 329:1521–1523.

Nagel, L., and D. Schluter. 1998. Body size, natural selection, and speciation in sticklebacks. Evolution 52:209–218.

Peichel, C. L., K. S. Nereng, K. A. Ohgi, B. L. Cole, P. F. Colosimo, C. A. Buerklet, D. Schluter, and D. M. Kingsley. 2001. The genetic architecture of divergence between threespine stickleback species. Nature 414:901–905.

Rennison, D. J., S. M. Rudman, and D. Schluter. 2019. Genetics of adaptation: Experimental test of a biotic mechanism driving divergence in traits and genes. Evolution Letters 3:513–520.

Rockman, M. V. 2012. The QTN program and the alleles that matter for evolution: All that’s gold does not glitter. Evolution 66:1–17.

Rogers, S. M., P. Tamkee, B. Summers, S. Balabahadra, M. Marks, D. M. Kingsley, and D. Schluter. 2012. Genetic signature of adaptive peak shift in threespine stickleback. Evolution 66:2439–2450.

Rundle, H. D. 2002. A test of ecologically dependent postmating isolation between sympatric sticklebacks. Evolution 56:322–9.

Rundle, H. D., L. Nagel, J. Boughman, D. Schluter, and J. Wenrick Boughman. 2000. Natural Selection and Parallel Speciation in Sympatric Sticklebacks. Science 287:306–308.

Rundle, H. D., and D. Schluter. 1998. Reinforcement of stickleback mate preferences: sympatry breeds contempt. Evolution 52:200–208.

Schemske, D. W., and H. D. Bradshaw. 2002. Pollinator preference and the evolution of floral traits in monkeyflowers (Mimulus). Proceedings of the National Academy of Sciences.

Schluter, D. 1995. Adaptive radiation in sticklebacks: Trade-offs in feeding performance and growth. Ecology 76:82–90.

Schluter, D. 2000. The Ecology of Adaptive Radiation. Oxford University Press, New York.

Schluter, D., K. B. Marchinko, M. E. Arnegard, H. Zhang, S. D. Brady, F. C. Jones, M. A. Bell, and D. M. Kingsley. 2021. Fitness maps to a large-effect locus in introduced stickleback populations. Proceedings of the National Academy of Sciences 118:e1914889118.

Schluter, D., and J. D. McPhail. 1992. Ecological Character Displacement and Speciation in Sticklebacks. The American Naturalist 140:85–108.

Schumer, M., and Y. Brandvain. 2016. Determining epistatic selection in admixed populations. Molecular Ecology 25:2577–2591.

Schumer, M., R. Cui, D. L. Powell, R. Dresner, G. G. Rosenthal, and P. Andolfatto. 2014. High-resolution mapping reveals hundreds of genetic incompatibilities in hybridizing fish species. eLife 3:e02535.

Simon, A., N. Bierne, and J. J. Welch. 2018. Coadapted genomes and selection on hybrids: Fisher’s geometric model explains a variety of empirical patterns. Evolution Letters 2:472–498.

Thompson, K. A. 2020. Experimental hybridization studies suggest that pleiotropic alleles commonly underlie adaptive divergence between natural populations. American Naturalist 196:E16–E22.

Thompson, K. A., M. Urquhart-Cronish, K. D. Whitney, L. H. Rieseberg, and D. Schluter. 2021. Patterns, predictors, and consequences of dominance in hybrids. The American Naturalist In press.

Vamosi, S. M., T. Hatfield, and D. Schluter. 2000. A test of ecological selection against young-of-the-year hybrids of sympatric sticklebacks. Journal of Fish Biology 57:109–121.

Van Ooijen, J., and R. Voorrips. 2001. Joinmap (R) version 3.0: Software for the calculation of genetic linkage maps.

Wang, R. J., M. A. White, and B. A. Payseur. 2015. The pace of hybrid incompatibility evolution in house mice. Genetics 201:229–242.

Zhang, Z., D. P. Bendixsen, T. Janzen, A. W. Nolte, D. Greig, and R. Stelkens. 2020. Recombining Your Way out of Trouble: The Genetic Architecture of Hybrid Fitness under Environmental Stress. Molecular Biology and Evolution 37:167–182.

